# Structure of the light harvesting 2 complex reveals two carotenoid energy transfer pathways in a photosynthetic bacterium

**DOI:** 10.1101/2020.10.21.349431

**Authors:** Alastair T. Gardiner, Katerina Naydenova, Pablo Castro-Hartmann, Tu C. Nguyen-Phan, Christopher J. Russo, Kasim Sader, C. Neil Hunter, Richard J. Cogdell, Pu Qian

## Abstract

We report the 2.4 Å resolution structure of the light harvesting 2 complex (LH2) from *Marichromatium (Mch.) purpuratum* determined by electron cryo-microscopy. The structure contains a heptameric ring that is unique among all known LH2 structures, explaining the unusual spectroscopic properties of this bacterial antenna complex. Two sets of distinct carotenoids are identified in the structure, and a network of energy transfer pathways from the carotenoids to bacteriochlorophyll *a* molecules is shown. The geometry imposed by the heptameric ring controls the resonant coupling of the long wavelength energy absorption band. Together, these details reveal key aspects of the assembly and oligomeric form of purple bacterial LH2 complexes that were previously inaccessible by any technique.

**One Sentence Summary:** The structure of a heptameric LH2 antenna complex reveals new energy transfer pathways and the basis for assembling LH rings.

---

In purple bacterial photosynthesis, light energy absorbed by LH2 is transferred via the light harvesting 1 complex to the reaction center complex, where it is trapped by the primary charge separation reactions. A detailed understanding of the molecular mechanisms of these reactions is firmly underpinned by knowledge gained from the structures of reaction center and light-harvesting complexes^1–8^. In this regard, the structures of the membrane-bound pigment-protein complexes from purple photosynthetic bacteria have been particularly influential. The most abundant of these is the LH2 antenna complex. All such complexes are formed by the circular oligomerization of dimer building blocks consisting of pairs of low molecular weight, hydrophobic apoproteins to which bacteriochlorophyll (BChl) and carotenoid (Car) molecules are non-covalently bound. The principles underlying the assembly, oligomerization, ring size and absorption properties of many natural variants of LH2 complexes are required for future designs of genetically modified and synthetic light absorbers, and therefore structural details are needed. Yet X-ray crystallographic studies have generally encountered difficulties with weak protein-protein contacts, lattice disorder and poor diffraction. High resolution single particle electron cryo-microscopy (cryo-EM) structures, required to correlate fine structural details with spectroscopic analyses, have until now been hampered by the small (<120 kDa) sizes of LH2 complexes and the variable amounts of radiation damage incorporated into the final reconstructed maps. We used a recently developed specimen support that allows imaging free of specimen movement and radiation damage artefacts^9^, to determine the 2.4 Å resolution structure of the LH2 complex from the marine purple bacterium *Mch. purpuratum*. We show how the circular packing of seven α/β subunits creates binding sites for two populations of carotenoid pigments, and how this organization determines the near-infra-red (NIR) absorption of excitonically coupled BChl *a* molecules.

Briefly, 17,338 movie files of the cryo-EM data were collected and 3.05 M particles were picked for data processing. Fig. 1A-C shows three views of the cryo-EM density map of the *Mch. purpuratum* LH2 complex at 2.4 Å resolution, which is a novel, circular LH2 structure formed of seven α/β apoprotein heterodimer subunits. Seven α-apoprotein α-helices form an inner ring of 25.4 Å diameter and the seven β-apoprotein α-helices form an outer ring of 55.0 Å diameter. The complex is surrounded by a 10 Å wide belt of disordered detergent molecules and the central hole has a diameter of 16 Å. Viewed from the side, at right angles to the long axes of the helices, the profile of the complex is strikingly trapezoidal, being broader on the cytoplasmic side. This feature is significantly enhanced compared with the previous X-ray structures of LH2 complexes.^3,4,10^.

**Fig. 1.**
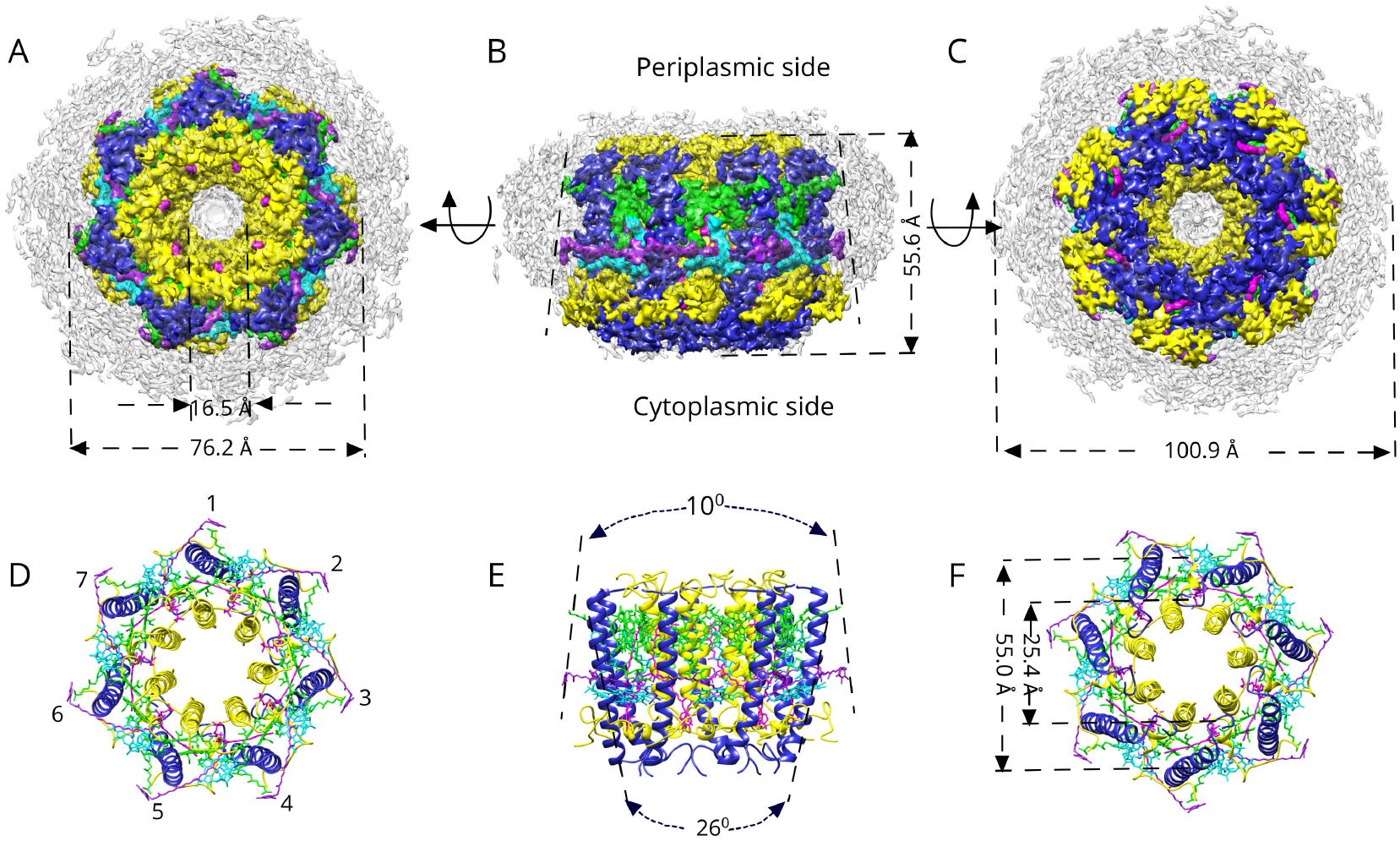
Cryo-EM structure of the LH2 from *Mch. purpuratum*. (**A-C**) The cryoEM density map of the LH2 complex is colored according to the color code below, and views from the cytoplasmic side of the membrane (**A**), in the membrane plane (**B**), and from the periplasmic side of the membrane (**C**) are shown. (**D-F**) Ribbon model views corresponding to the views and coloring in (**A-C**.) are shown. The α/β–heterodimers are numbered in (**D**). Viewed as in **B**, showing the marked tapering of trans-membrane helices and the reversed 10° taper and trapedoizal shape of the whole LH2 complex in (**E**). The diameters of the inner α and outer β rings of the LH2 in (**F**) are measured from the centers of the respective helices, midway through the transmembrane region of the complex. Color code: α–polypeptide, *yellow*; β-polypeptide, *medium blue*; B828, *green*; B800, *cyan*; Car1, *red*; Car2, *purple*; detergent belt, *grey*.

Previous mass spectrometry analyses^11^ found three homologous α-polypeptides and three homologous β-polypeptides in the purified *Mch. purpuratum* LH2 complex (Fig. S1, Fig. S2). The exact distribution of the different but highly homologous α/β-pairs in a single LH2 molecule is unknown. A 2.8 Å resolution reconstruction of the LH2 without symmetry imposed (Fig. S3) shows a pseudo-C7 symmetry up to this resolution (Fig. S4B). This indicates that either the different α/β-pairs are randomly distributed in the heptameric LH2 ring, and/or that the different α/β-pairs have very similar structures. Comparison of the amino acid sequences of the three different α-polypeptides (Fig. S1) shows that they differ in their N–termini (residues 1-15) and at residue α-47 (MML), which lies in the helical region. The relatively low resolution and high atomic *B*–factors in the N–terminal region of the α-polypeptide in the structure (Fig. S4D) are probably due to mixed densities from the three different sequences and the intrinsic flexibility of the loop, which is also seen in the C–terminal region. Only two residues differ in the central helical regions of the three types of β-polypeptide (positions β-14 (AAE) and β-43 (VII), and the N– and C–terminal loops are much shorter than for the α-polypeptides. Since none of these differences could be resolved, we imposed C7 symmetry on the three-dimensional reconstruction, producing an LH2 structure close to the native LH2 from *Mch. purpuratum.*

The complete α-apoprotein can be traced in the structure apart from the N-terminal and C-terminal amino acids, and the β-apoprotein can be fully traced apart from the first four N-terminal amino acids. Both apoproteins have a single membrane-spanning α-helix, with their N-termini on the cytoplasmic side of the complex and their C-termini on the periplasmic side (Fig. 2A). There are two strong H-bonds between adjacent heterodimers, between α-Ser 63 and β-Phe 48 on the periplasmic side, and between β-Gln 15 and β-Asn 7 on the cytoplasmic side (Fig. 2B). Each α/β–heterodimer binds, non-covalently, three molecules of BChl *a* and two molecules of the carotenoid (Car), okenone. Two of the BChl *a* molecules form a strongly interacting dimer with their Mg^2+^ atoms liganded by two His residues, α-His 49 and β-His 38. The distance between these two Mg^2+^ atoms is 9.6 Å. By analogy with previous LH2 structures, the 828 nm absorption band is assigned to these two BChl *a* molecules. Their orientations are fixed by a set of H-bonds between the C-17^3^ carbonyl group and β-Trp 30 for the α–BChl *a* molecules, and between the C-13^2^ carbonyl group of the β-BChls *a* and α-Gln 44. The third monomeric BChl *a* molecule is located on the cytoplasmic side of the complex; its Mg^2+^ atom is liganded to α–Asp 21 and there is an H-bond between its C-3^1^ carbonyl group and β-His 25 from the β-apoprotein in the next heterodimer (Fig. 2B). This BChl *a* molecule can be assigned to the 800 nm absorption band. One molecule of the carotenoid okenone (Car1) has a twisted, all-*trans* configuration (Fig. S5) and runs approximately parallel to the heterodimer α helices across most of the length of the complex. The region of its conjugated C=C double bonds comes into close contact (3.6 Å) with the bacteriochlorin ring of the B828 α–BChl *a* molecule and runs parallel to the Q_y_ transition dipole moment of that bacteriochlorin ring (Fig. 2C). A second okenone molecule (Car2) adopts a 9-*cis* configuration (Fig. S5) and lies in the plane of the membrane, at right angles to the heterodimer α-helices. Its region of C=C conjugated double bonds run parallel to the Q_x_ transition dipole moment of the B800 BChl *a* at a closest distance of 3.2 Å (Fig. 2C, 2D). The presence of this Car2 in every heterodimer subunit and the fact that it adopts a 9-*cis* configuration are both unique features of this LH2 complex. All inter- / intra-subunit protein-protein and protein-pigment interactions are summarized in Fig. 2E.

**Fig. 2.**
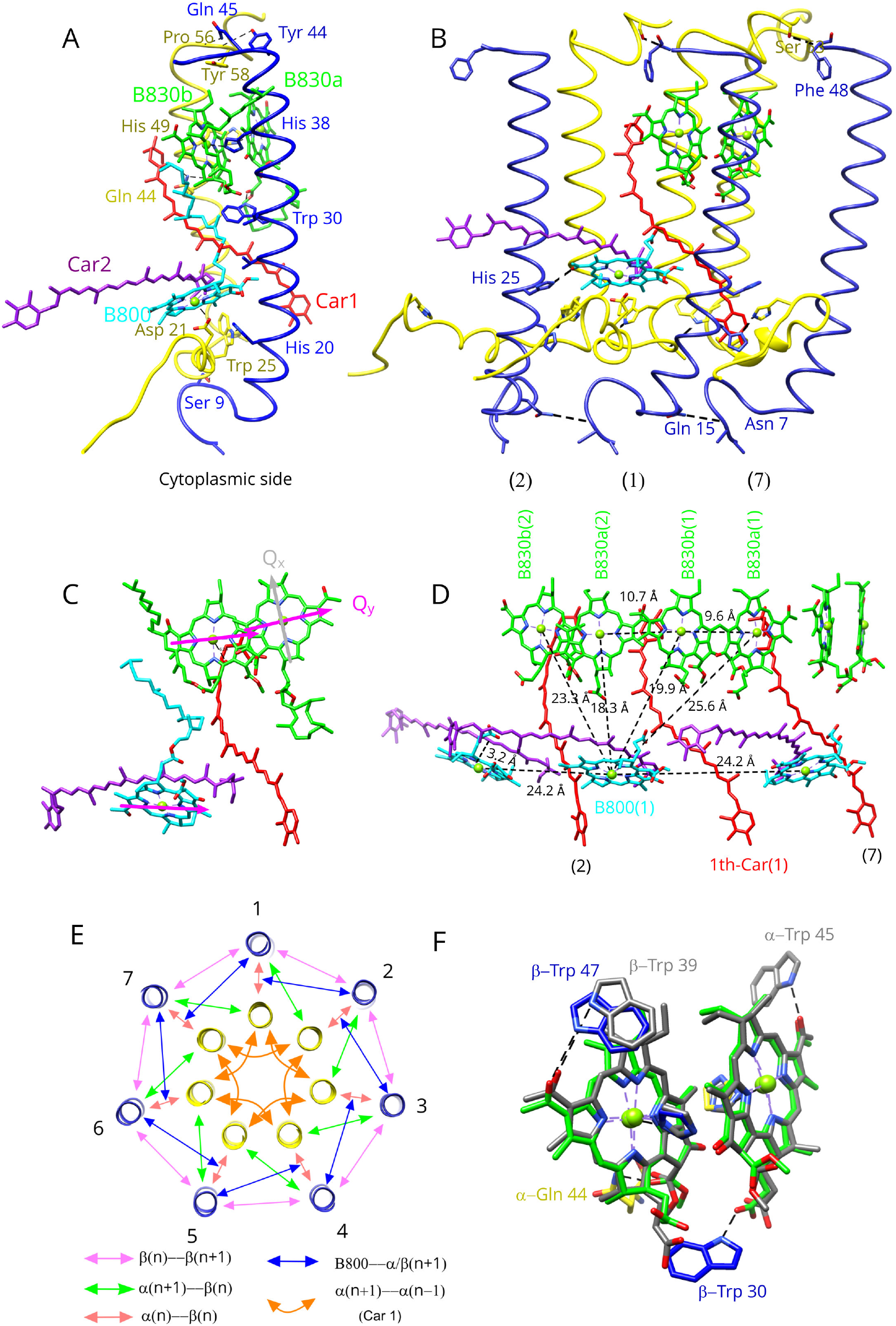
Protein-protein and protein-pigment interactions in the LH2 from *Mch. purpuratum*. (**A**) The molecular model of one subunit of the LH2 complex is shown. Only the residues involved in protein-protein and protein-pigment interactions are labeled for clarity. (**B**) Three subunits are presented to show the interactions between them, but only one unit includes pigments for clarity. The phytol tails of the BChls are omitted. (**C**) All of the pigments within one subunit are shown and the BChl *a* Qx and Qy transitions are indicated in *grey* and *magenta*, respectively. (**D**) The pigment arrangement in three consecutive subunits of the LH2 determines the distances between Mg atoms of the BChls. The BChl *a* tails are omitted. (**E**) All protein-protein and protein-pigment interactions in the LH2 complex are summarized schematically. (**F**) The B828 pair from *Mch. purpuratum* (*green*) and the B858 pair from *Rbl. acidophilus* (*grey*) are superimposed for comparison. The color code is the same as in Fig.1: α–polypeptide, *yellow*; β-polypeptide, *medium blue*; B828, *green*; B800, *cyan*; Car1, *red*; Car2, *purple*.

**Fig. 3.**
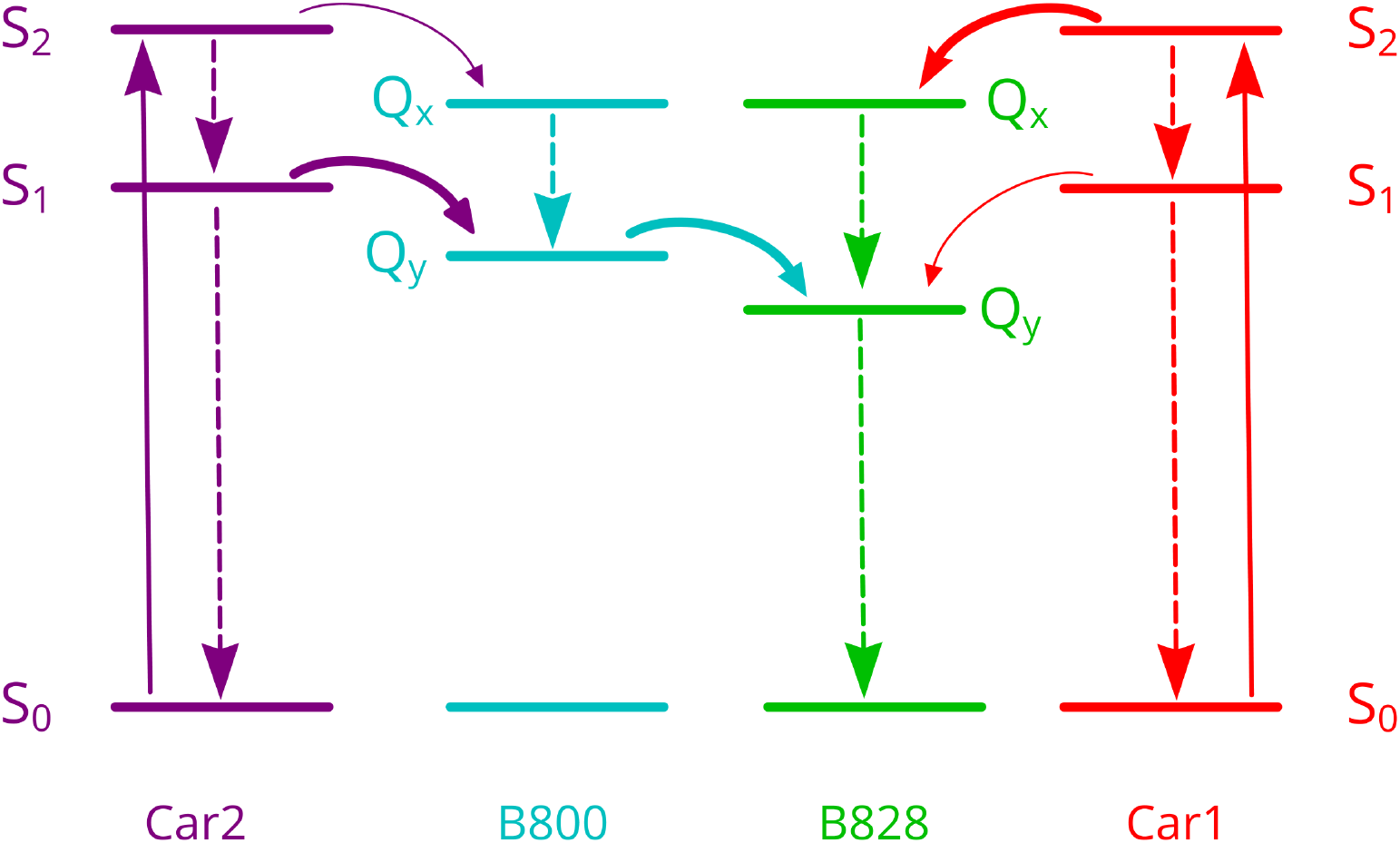
Schematic diagram of energy levels and energy transfer pathway from Car to BChl *a* in the LH2 from *Mch. purpuratum.* Solid curved arrows represent energy transfer pathways, and dashed arrow lines denote internal conversions. Color code is same as in Fig. 1. Less important, possible energy transfer pathways from Car1 to B800 and Car2 to B828 are omitted for brevity.

Three basic factors determine the position of the long wavelength Q_y_ absorption band arising from the ring of tightly coupled BChl *a* molecules in purple bacterial LH2 complexes ^7,12,13^: (i) the ‘site’ energy of the individual BChl *a* molecules in their respective binding sites, (ii) the coupling between BChl *a* molecules within their α/β-subunits to form a dimer, and (iii) the interactions between dimers in adjacent α/β-subunits around the LH2 ring. In the case of LH2 from *Rhodoblastus (Rbl.) acidophilus* (formerly *Rhodopseudomonas (Rps.) acidophila*) the strongly coupled ring of BChl *a* molecules have their Q_y_ absorption band at 858 nm, whereas in the LH2 complex from *Mch. purpuratum* this absorption band is blue-shifted to 828 nm (Fig. S6) The organization, orientation, and H-bonding of the bacteriochlorin rings of both BChl *a* dimers are very similar between these two species (Fig. 2E), in terms of in-plane orientation of their carbonyl groups relative to the plane of the bacteriochlorin rings, and their Mg^2+^-Mg^2+^ distances, which are 9.6 Å in the case of *Mch. purpuratum* and 9.4 Å in the case of *Rbl. acidophilus*. Although these similarities could suggest similar dimer exciton bands in these LH2 complexes, there is a large difference in the Mg^2+^-Mg^2+^ distances between one of the BChl *a* molecules in a heterodimer subunit and the nearest BChl *a* molecule in the next heterodimer subunit in the LH2 ring. In the case of *Rbl. acidophilus* this distance is 8.8 Å, whereas for *Mch. purpuratum* it is 10.7 Å (Fig. 2D). This difference, as well as the increased angle between dimers imposed by the 7-mer ring relative to a 9-mer, lowers the strength of dimer-dimer exciton coupling, which causes the blue-shift of the B850 band in *Rbl. acidophilus* to B828 in *Mch. purpuratum*.

Carotenoids have a strong visible absorption band arising from the optically allowed transition from the ground state, S_0_, to the 2^nd^ excited singlet state, S_2_^14,15^. The lower lying, symmetry-forbidden 1^st^ excited singlet state, S_1_, is populated by ultrafast internal conversion from the S_2_ state^16,17^. In most purple photosynthetic bacteria, carotenoids function as accessory light harvesting pigments and can transfer absorbed solar energy to the BChl molecules thereby making it available to drive photosynthesis^18^. A previous study of ultrafast energy transfer within isolated LH2 complexes from *Mch. purpuratum* showed that the efficiency of energy transfer from okenone to BChl *a* is 95+/− 5%^19^. This Car to BChl *a* energy transfer can take place either from the Car S_2_ state alone or from both the S_2_ and S_1_ states, depending on the type of carotenoid involved. One okenone transfers its excitation energy to B828 in <200 fs from S_2_ and in 3.8 ps from S_1_, and the other transfers its excitation energy to B800 in about 0.5 ps. The results of this study could only be explained by assuming the presence of two separate pools of okenone with different orientations of their transition dipole moments relative to BChl *a* molecules. The structure of the LH2 complex from *Mch. purpuratum* described here is the first direct observation of these two populations of different carotenoids, which serve structurally and spectroscopically distinct BChl *a* molecules, and accounts for the spectroscopic observation, which has lain unexplained for more than 24 years. The 3.6 Å distance (Fig 2B) is consistent with the rapid (<200 fs) energy transfer from Car1 to the nearest B828 pigment, and Car2 is precisely positioned, with a distance of 3.2 Å, to transfer its excitation energy to B800. The pigment organization and their configurations in the LH2 complex are shown in Fig. S7 and Fig. S8.

Trying to understand the factors that control the ring size of purple bacterial LH2 complexes is a long-standing problem. Fig. 4 compares the details of the interactions of the α/β heterodimers in the structures of the LH2 complexes from *Rbl. acidophilus* (9-mer) and *Mch. purpuratum* (7-mer). This comparison provides a clear explanation for their different oligomeric sizes.

**Fig. 4.**
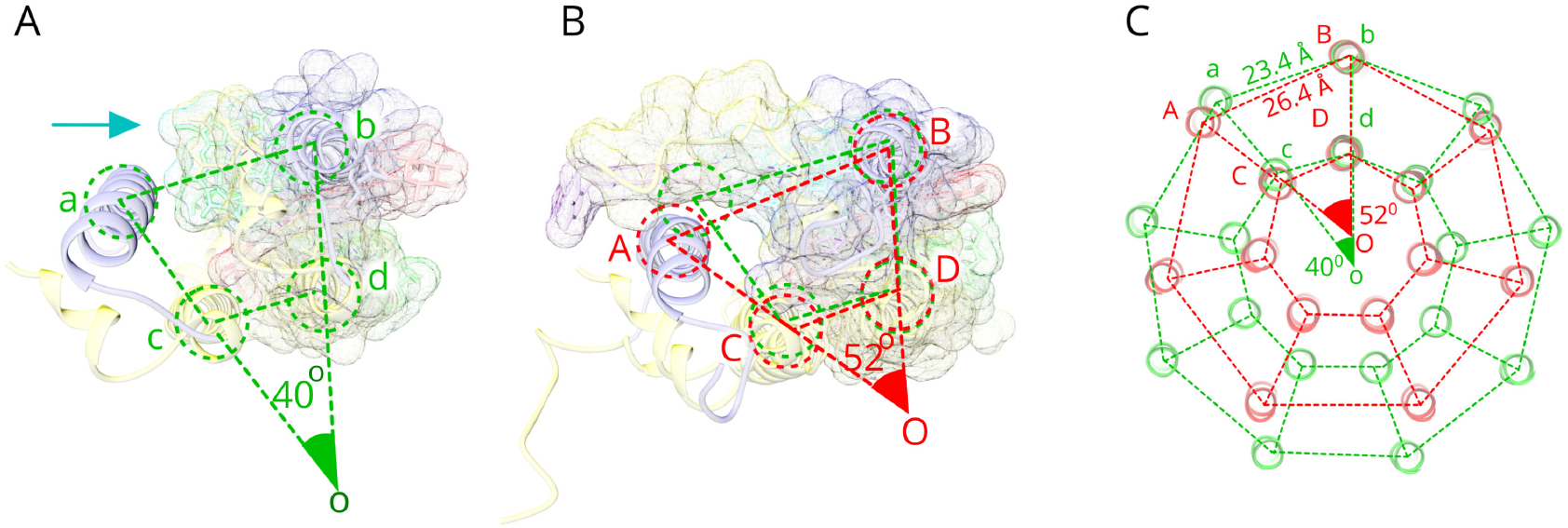
Assembly mechanism of LH2 complexes from *Rps. acidophila* and *Mch. purpuratum*. A pair LH2 subunits from *Rbl. Acidophilus* (**A**) and *Mch. Purpuratum* (**B**), viewed from the cytoplasmic side of the plasma membrane. An *arrow* points to N-terminal domain of the α–polypetide in (**A**). The dashed *green/red* circles represent the positions of trans-membrane helices. The wider A-B spacing (*red*) is compatible with the second okenone molecule, Car2 (*purple*), and a longer N-terminal region of the α-polypeptide (*yellow*), whereas the shorter a-b spacing is not. **(C**) The schematic representation of LH2 complexes from *Rbl. acidophilus* (*green*) and *Mch. purpuratum* (*red*), with their segments in as in (**A**) and (**B**) illustrates their two different patterns of oligomerization.

In the case of the 9-mer LH2 complex from *Rbl. acidophilus*, a line drawn from the center of a pair of β-helices (ab), followed by a pair of lines from the center of each β helix to the center of their heterodimer α apoprotein helix (ac and bd) form a segment that has a 40° angle between these two lines (∠aob = 360°/9 = 40°) (Fig. 4A). In the case of the 7-mer LH2 complex from *Mch. purpuratum*, each α/β heterodimer relates to the adjacent segment by a ~52° rotation (∠AOB=360°/7=52.14°) (Fig. 4B). Figure 4C illustrates how these two patterns of oligomerization are manifested at the level of intact LH2 complexes. This comparison provides new insights into way that LH2 α/β heterodimers fit together, and in particular the basis for the assembly of a 7-mer ring in the *Mch. purpuratum* LH2 antenna complex. Firstly, as shown in Fig. 4B, the longer N-terminal region of the α apoprotein and the second okenone molecule, Car2, protrudes from the α/β heterodimer in a way that would induce strong steric clashes if the segment angle was reduced to 40° as in the case of 9-mer *Rbl. acidophilus*. A wider segment angle, namely ~52°, is needed to circumvent these potential clashes necessitating a 7-mer complex. Therefore, as a direct consequence of this work, we can clearly visualize the main factors involved in controlling this segment angle and, thereby, the oligomeric size of the LH2 ring.

The extended N-terminal domain for LH2α modifies the packing of α/β heterodimers, widening their spacing by 12° and creating space for binding an extra carotenoid (Fig. 4B). This packing of neighboring α/β–apoprotein dimers in the holo-LH2 structure also determines the overall oligomeric ring size, the number of excitonically coupled BChl *a* molecules, the extent of coupling and the position of the long wavelength absorption band. Thus, the structure of the 7-mer LH2 complex from *Mch. purpuratum* provides a detailed structural understanding of how the oligomeric ring size of LH2 complexes is controlled, and explains its previously determined spectroscopic and biochemical properties.

Many species of purple bacteria have multiple *puc* genes encoding LH2 apoproteins suggesting that the heterologous LH2 complexes, made from different α/β-apoprotein pairs, are much more common than previously appreciated^20,21^. Our structural analysis of such a heterologous complex, from *Mch. purpuratum*, by cryo-EM circumvents the difficulties encountered over many years in X-ray crystallography, where heterologous LH2 complexes weaken protein-protein contacts, leading to long-range internal disorder within the lattice and poor diffraction. Technical developments in cryo-EM, including direct electron detectors, high-speed data acquisition, movement-free specimen supports, and image processing algorithms circumventing radiation damage, have enabled us to determine the structure of this small, heterologous LH2 complex to a resolution sufficient to identify and measure the critical features of the pigments and understand the energy transfer pathway.

## Acknowledgments

K.N. and C.J.R. thank J. Grimmett and T. Darling of the LMB Scientific Computing for technical assistance with computation.

## Author contributions

P.Q, C.N.H., R.J.C, C.J.R and K.S. conceptualized the project. A.T.G., K.N, P.C.-H., T.C.N.-P. and P.Q. performed experiments. P.Q., R.J.C, C.J.R., K.N., A.T.G. and C.N.H. wrote the manuscript. All authors edited and revised the manuscript.

## Data and materials availability

Atomic model and 3D cryo-EM map are available on wwPDB and EMDB with access codes of 6ZXA and EMD-11516.

## References and Notes

1 Deisenhofer, J., Epp, O., Miki, K., Huber, R. & Michel, H. X-Ray structure analysis of a membrane protein complex electron density map at 3Å resolution and a model of the chromophores of the photosynthetic reaction center from *Rhodopseudomonas viridis*. Journal of Molecular Biology 180, 385–398 (1984).

2 Allen, J. P. et al. Structural homology of reaction centers from *Rhodopseudomonas sphaeroides* and *Rhodopseudomonas viridis* as determined by X-ray diffraction. Proc.Natl.Acad.Sci.U.S.A 83, 8589–8593 (1986).

3 McDermott, G. et al. Crystal structure of an integral membrane light-harvesting complex from photosynthetic bacteria. Nature 374, 517–521 (1995).

4 Koepke, J. et al. The crystal structure of the light-harvesting complex II (B800-B850) from *Rhodospirillum molischianum*. Structure 4, 581–597 (1996).

5 Yu, L. J., Suga, M., Wang-Otomo, Z. Y. & Shen, J. R. Structure of photosynthetic LH1-RC supercomplex at 1.9 Å resolution. Nature 556, 209–213 (2018).

6 Umena, Y., Kawakami, K., Shen, J. R. & Kamiya, N. Crystal structure of oxygen-evolving photosystem II at a resolution of 1.9 Å. Nature 473, 55–60, doi:10.1038/nature09913 (2011).

7 Qian, P., Siebert, C. A., Wang, P. Y., Canniffe, D. P. & Hunter, C. N. Cryo-EM structure of the *Blastochloris viridis* LH1-RC complex at 2.9 angstrom. Nature 556, 203–208 (2018).

8 Liu, Z. F. et al. Crystal structure of spinach major light-harvesting complex at 2.72 angstrom resolution. Nature 428, 287–292 (2004).

9 Naydenova, K., Jia, P. & Russo, C. J. Electron cryomicroscopy with sub-Angrtrom specimen movement. Science 370, 223–226 (2020).

10 McLuskey, K., Prince, S. M., Cogdell, R. J. & Isaacs, N. W. The crystallographic structure of the B800-820 LH3 light-harvesting complex from the purple bacteria *Rhodopseudomonas acidophila* strain 7050. Biochemistry 40, 8783–8789 (2001).

11 Cranston, L. J. Structural and functional studies of LH2 complexes having unusual spectroscopic properties. PhD thesis, University of Glasgow, (2017).

12 Ma, F., Yu, L. J., Wang-Otomo, Z. Y. & van Grondelle, R. The origin of the unusual Q(y) red shift in LH1-RC complexes from purple bacteria *Thermochromatium tepidum* as revealed by Stark absorption spectroscopy. Biochim. Biophys. Acta 1847, 1479–1486 (2015).

13 Cogdell, R. J., Howard, T. D. & Isaacs, N. W. Structural factors which control the position of the Qy absorption band of bacteriochlorophyll *a* in purple bacterial antenna complexes Photosynth. Res. 74, 135–141 (2002).

14 Schulten, K. & Karplus, M. On the origin of a low-lying forbidden transition in polyenes and related molecules. Chem. Phys. Lett., 305–309 (1972).

15 Hudson, B. S. & Kohler, B. E. A low-lying weak transition in the polyene α,ω-diphenyloctatetraene. Chem. Phys. Lett. 14, 299–304 (1972).

16 Shreve, A. P., Trautman, J. K., Owens, T. G. & Albrecht, A. C. Determination of the S2 lifetime of β-carotene. Chem. Phys. Lett. 178, 89–96 (1991).

17 Macpherson, A. N. & Gillbro, T. Solvent dependence of the ultrafast S2–S1 internal conversion rate of β-carotene. J. Phys. Chem. 102, 5049–5058 (1998).

18 Cogdell, R. J. et al. How carotenoids protect bacterial photosynthesis. Philos. T. R. Soc. B 355, 1345–1349 (2000).

19 Andersson, P. O., Cogdell, R. J. & Gillbro, T. Femtosecond dynamics of carotenoid-to-bacteriochlorophyll a energy transfer in the light-harvesting antenna complexes from the purple bacterium *Chromatium purpuratum* Chem. Phys. 210, 195–217 (1996).

20 Brotosudarmo, T. H. P. et al. Single-Molecule Spectroscopy Reveals that Individual Low-Light LH2 Complexes from *Rhodopseudomonas palustris* 2.1.6. Have a Heterogeneous Polypeptide Composition. Biophys. J. 97, 1491–1450 (2009).

21 Gardiner, A. T., Niedzwiedzki, D. M. & Cogdell, R. J. Adaptation of *Rhodopseudomonas acidophila* strain 7050 to growth at different light intensities: what are the benefits to changing the type of LH2? Faraday Discuss 207, 471–489 (2018).

22 Imhoff, J. F. & Truper, H. G. Chromatium purpuratum, sp. nov., a new species of the Chromatiaceae. Zentralblatt für Bakteriologie: I. Abt. Originale C: Allgemeine, angewandte und ökologische Mikrobiologie 1, 61–69 (1980).

23 Bellare, J. R., David, H. T., Scriven, L. E. & Talmon, Y. Controlled Environment Vitrification System: An Improved Sample Preparation Technique. J. Electron Micr. Tech. 10, 87–111 (1988).

24 Russo, C. J., Scotcher, S. & Kyte, M. A precision cryostat design for manual and semi-automated cryo-plunge instruments. Rev. Sci. Instr. 87, 114302 (2016).

25 Sader, K., Matadeen, R., Hartmann, P. C., Halsan, T. & Schlichten, C. Industrial cryo-EM facility setup and management. Acta Cryst. D 76, 313–325 (2020).

26 Zivanov, J. et al. New tools for automated high-resolution cryo-EM structure determination in RELION-3. Elife 7, e42166, doi:10.7554/eLife.42166 (2018).

27 Zhang, K. Gctf: Real-time CTF determination and correction. J. Struct. Biol. 193, 1–12 (2016).

28 Grant, T., Rohou, A. & Grigorieff, N. cisTEM, user-friendly software for single-particle image processing. Elife 7, e35383, doi:10.7554/eLife.35383 (2018).

29 Li, X. M. et al. Electron counting and beam-induced motion correction enable near-atomic-resolution single-particle cryo-EM. Nature methods 10, 584–+ (2013).

30 Rohou, A. & Grigorieff, N. CTFFIND4: Fast and accurate defocus estimation from electron micrographs. J. Struct. Biol. 192, 216–221 (2015).

31 Zivanov, J., Nakane, T. & Scheres, S. H. W. A Bayesian approach to beam-induced motion correction in cryo-EM single-particle analysis. Iucrj 6, 5–17 (2019).

32 Pettersen, E. F. et al. UCSF Chimera--a visualization system for exploratory research and analysis. J. Comput. Chem. 25, 1605–1612, doi:10.1002/jcc.20084 (2004).

33 Emsley, P. & Cowtan, K. Coot: model-building tools for molecular graphics. Acta Cryst. D 60, 2126–2132 (2004).

34 Murshudov, G. N. et al. REFMAC5 for the refinement of macromolecular crystal structures Acta Cryst. D 67, 355–367 (2011).

35 Liebschner, D. et al. Macromolecular structure determination using X-rays, neutrons and electrons: recent developments in Phenix. Acta Cryst. D75, 861–877 (2019).

36 Rosenthal, P. B. & Henderson, R. Optimal determination of particle orientation, absolute hand, and contrast loss in single-particle electron microscopy. J. Mol. Biol. 333, 721–745 (2003).

37 Naydenova, K. & Russo, C. J. Measuring the effects of particle orientation to improve the efficiency of electron cryomicroscopy. Nat Commun 8, 1–5, doi:10.1038/s41467-017-00782-3 (2017).

38 Kucukelbir, A., Sigworth, F. J. & Tagare, H. D. Quantifying the local resolution of cryo-EM density maps. Nature methods 11, 63–67 (2014).

