## Supplemental materials for "Structure of the light harvesting 2 complex reveals two carotenoid energy transfer pathways in a photosynthetic bacterium"

### Protein purification

*Mch. purpuratum* strain BN5500 (also designated as DSM1591 or 984) was grown anaerobically in the light in Pfennig's medium[22], with incandescent bulbs at a light intensity of ~80 µmol $m^{-2}$ $s^{-1}$ at 30 °C. Harvested cells were washed once with 20 mM MES, 100 mM KCl, pH 6.8 and the pellet flash-frozen until required. The cell pellet was defrosted and re-suspended in 20 mM Tris.HCl buffer pH 8.0, with a few grains of DNase and $MgCl_2$ added, homogenized and then disrupted by passing twice through a French Press cell at ~15,000 psi. Unbroken cells and debris were removed by a low-speed spin (3,000 g, 10 min, 4 °C) and the chromatophore membranes pelleted by ultracentrifugation (180, 000 g, 90 min, 4 °C). The chromatophores were re-suspended in 20 mM Tris.HCl pH 8.0 buffer to an optical density (OD) = 50 $cm^{-1}$ at the NIR absorbance maximum (~828 nm). The sample was solubilised by the addition of N,N Dimethyldodecylamine N-oxide (LDAO) to 1.0% (v/v), with stirring for 1 min and the mixture was immediately loaded on to a pre-equilibrated glass gravity Q-Sepharose (GE Healthcare) column. The sample was washed with copious amounts of 0.02 % n-dodecyl-β-D-maltopyranoside (DDM) in 20 mm Tris.HCl pH 8.0 (called TD buffer) and then the LH2 was eluted with increasing concentrations of NaCl in TD buffer. The LH2-rich fractions were assayed, pooled and dialysed overnight in TD buffer to remove the NaCl. Due to the huge amount of free pigment involved, the process was repeated the following day by loading the dialysed sample on to a fresh Q-Sepharose column. The eluted LH2 was then assayed, pooled and concentrated prior to passage down a Superdex G200 gel filtration column (GE Healthcare) with TD buffer. Fractions having an $A_{828}$ /$A_{280}$ ratio of 2.2 or higher were concentrated to an absorbance at 828 nm = 100 $cm^{-1}$ for cryo-EM grid preparation.

### Cryo-EM data collection

Two different grids were used for cryo-EM specimen preparation. Initially, a QuantiFoil R1.2/1.3 400 mesh Cu grid, was glow discharged for 30 seconds (easiGlow). The grid was plunge frozen into liquid ethane using a FEI Vitrobot MK IV, equilibrated to 100% humidity at 4 $^{o}$C. A sample volume of 3µl was applied to the grid and it was blotted for 2.5 seconds before freezing. For the final data collection, a HexAufiol grid[9], manufactured in house at the LMB, was used. The grid was plasma cleaned under a mixed atmosphere ($O_2$:Ar = 1:9) for 60 sec in a plasma chamber for 60 seconds (Fischione 1070). It was vitrified using a manual plunger of the Talmon type[23] in a 4 $^{o}$C cold room, and an ethane cryostat[24] held at 93K. 3 μl protein solution was added applied to the foil side of the cleaned grid, and manually blotted for 11 seconds with filter paper (Whatman). All were stored in liquid nitrogen before use. Data were collected on a Thermofisher Titan Krios G3i electron cryo-microscope equipped with a Falcon 4 direct electron detector at the Cambridge Pharmaceutical Cryo-EM Consortium[25]. The microscope was operated at 300 kV with a nominal magnification of 120,000×, corresponding to 0.646 Å /pixel at the specimen level, calibrated using the Au (111) lattice reflections. The detector was operated in counting mode at a flux of 3.58 $e^{-}$/ $Å^2$/s. Each 12.18 s exposure was fractionated into 42 frames, resulting in an electron dose of 1.04 $e^{-}/Å^2$/frame. The defocus range was set to -0.8 to -2.4 μm. Automated data acquisition was performed in EPU 2.6 (ThermoFisher Scientific) with one exposure per hole in aberration-free image shift (AFIS) mode. In total, 7795 movies were collected from the QuantiFoil R1.2/1.3 grid, and 8935 movies - from the HexAuFoil grid (Fig. S9)

**Cryo-EM data processing**
The initial dataset collected from the QuantiFoil grid was processed in RELION 3.1. The movie stacks were motion-corrected within RELION[26] on 5 × 5 patches. The CTF parameters were determined using Gctf[27]. The particles were auto picked in cisTEM[28], and their coordinates imported in RELION for particle extraction using a box size of 270 × 270 pixels. A total of 1,723,876 particles were extracted, and subjected to 2D reference-free classification, then 1,337,902 particles were selected from good 2D classes. Reference free 2D classification showed that the LH2 from *Mch. purpuratum* is a heptamer with a similar architecture to other LH2 complexes. The initial heptamer model for 3D classification was built from an α/β subunit taken from the *LH2 of Phs. molischinum* (PDB 1LGH) using Chimera[28]. At this stage, C7 symmetry was imposed for 3D reconstructions. The best 3D class (3.98 Å), out of 4 classes, contained 867,046 (50.3%) particles. After multiple rounds of 3D refinement, anisotropic magnification, beam-tilt, trefoil, 4$^{th}$ order aberration, per-particle defocus, and per-micrograph astigmatism estimation, and particle movement tracking using Bayesian polishing with the default parameters in RELION, these 867,046 particles produced a 2.48 Å resolution map.

The dataset from the HexAuFoil grid was processed according to the previously proposed method for handling movement-free cryo-EM data[26]. The whole micrograph movement was corrected using MotionCorr[29]. The motion-corrected stacks we imported in Relion 3.1. A total of 1,330,145 particles were auto picked in cisTEM. The CTF correction was performed in CTFFIND4.1[30]. The particles were extracted into 270 × 270 pixel boxes, and subjected to 3D classification with a reference taken from previously determined map with 30 Å initial low pass filter applied. In total, 414,511 (31.2%) particles from the best 3D class (out of 4 classes) were selected. The selected particles were re-extracted using 512 x 512 box size for CTF refinement and Bayesian polishing[31]. No significant particle movement could be fit by the Bayesian polishing, as judged by the resolution of the reconstruction before and after this step. Similarly, no significant aberrations or anisotropic magnification were found. Per-particle defocus and/or astigmatism refinement did not yield any improvement compared to per-micrograph estimation of these parameters. We also tried separating the data into 149 optical groups based on image beam shift values, and estimating the optical aberrations for each group separately, but no significant variation between these groups was found. The final 3D reconstruction reached 2.38 Å resolution with C7 symmetry (2.76 Å without symmetry). Per-frame reconstructions were produced from particles extracted at their refined positions from individual frames of the aligned stacks. These reconstructions were used to extrapolate the structure factors, i.e. phases and amplitudes, in the 10 Å to 2.5 Å resolution range to zero electron dose, yielding the final map.

### Modeling and refinement

A α/β subunit polypeptide pair taken from the *LH2 complex of Phs. molischianum* (PDB 1LGH) was docked into the C7 symmetry imposed cryo-EM map as a template using Chimera[32] such that three BChl *a*s were roughly fitted with their corresponding densities. Mass spectroscopy (MALDI-TOF) of the *Mcr. purpuratum* LH2 revealed that three different α– and three different β–polypeptide are incorporated into the LH2 complex of *Mch. purpuratum*. They are distributed randomly, and with unknown stoichiometries, in the LH2 complex. Imposing C7 symmetry on the cryo-EM map of the LH2 from *Mch. purpuratum* complex during refinement mixed three different α– or β–polypeptides together, resulting in an averaged single α– and single β–polypeptide in the density map of the LH2 complex. In this case, the longest α– and β–amino acid sequences, i.e., α3 and β2 (Fig. S1) were selected for mutation of amino acids in the template using COOT[33]. Carotenoid lycopene in the template was replaced with an all-*trans* carotenoids okenone. The second carotenoid was fitted with 9-*cis* okenone confidently (Fig. S5, S6). Thus, a subunit of the LH2 from *Mch. purpuratum*, $\alpha_1\beta_1Car_2BChla_3$, was constructed. This subunit was then copied into the LH2 density map using the rigid body fitting in the Chimera, forming an atomic model of the heptameric LH2 complex. The model was real space refined in COOT[33]. A geometry-optimized model was then subjected to global refinement using Refmac5[34] and Phenix[35]. The refinement statistics are summarized in Table S1. The refined model and its cryo-EM map were deposited in the PDB and EMDB with codes of 6ZXA and EMD-11516.

LH2 α–polypeptides

| | | M.W. |
|---|---|---|
| 1. | MNCGKIWTVVNPAIGIPALLGSVTVIAILVHLAILSHTTWFPAYWCGGVKKAA | 5.60 |
| 2. | MNCGKIWTVVPPAFGLPLMLGAVAITALLVHAAVLTHTTWYAAFLCGGVKKAA | 5.50 |
| 3. | SNPKDDYKIWLVINPSTWLPVIWIVATVVAIAVHAAVLAAPGFNWIALGAAKSAAK | 5.94 |
| 4. | SNPKDDYKIWLVINPSTWLPVIWIVATVVAIAVHSFVLSVPGYNFLASAAAKTAAK | 6.06 |
| 5. | MTNGKIWLVVKPTVGVPLFLSAAVIASVVIHAAVLTTTTWLPAYYQGSAAVAAE | 5.60 |
| 6. | MQVPVMMGDPNAKLNHPEDDWKIWTVINPATWMVPFFGILFVQMWMIHSYALSLPGYGFKDSVRVAQPAA | 7.99 |
| 7. | MQVPVLLADKDVKLNHPEDDWKIWTVINPATWMVPFFGILFVQMWMIHSYALSLPGYGFKDSVRVAQPAA | 8.03 |
| 8. | MKVPVMMADESIATINHPEDDWKIWTVINPATWMVPFFGILFVQMWLIHSYALSLPGYGFKDSVRVAQPAA | 8.08 |
| | 1 10 20 30 40 50 60 70 | |

LH2 β–polypeptides

| | | |
|---|---|---|
| 1. | ATLTAEQSEELHKYVIDGTRVFLGLALVAHFLAFSATPWLH | 4.55 |
| 2. | AEVLTAEQAEELHKHVIDGTRVFLVIAAIAHFLAFTLTPWLH | 4.71 |
| 3. | AERSLSGLTEEEAIAVHDQFKTTFSAFIILAAVAHVLVWVWKPWF | 5.12 |

```
4.               AERSLSGLTEEEAVAVHDQFKTTFSAFIILAAVAHVLVWIWKPWF            5.12
5.          MTDDLNKVWPSGLTVAEAEEVHKQLILGTRVFGGMALIAHFLAAAATPWLG           5.45
6.                 ANLSGLTDAQAKEFHEHWKHGVWSWVMIASAVHVVTWVYQPWF            5.05
7.           ADPKAANLSGLTDAQAKEFHEHWKHGVWSWVMIASAVHVVTWIYQPWF            5.55
8.             AESKNLSGLSDEQAKEFHEHWKHGVWSWVMIASAVHVVTWIYQPWF            5.45
              1        10        20        30        40
```

```
1–Rps. acidophila 10050
2–Rps. acidophila 7050
3–Phs. molischinum DSM-119
4–Phs. molischinum DSM-120
5–Rba. sphaeroides 2.4.1
6–Mch. purpuratum  DSM-1591 1
7–Mch. purpuratun  DSM-1591 2
8–Mch. purpuratum  DSM-1591 3
```

Fig. S1 **Amino acid sequence alignments of α and β-polypeptides in LH2 complexes.** There are three α- and three β-polypeptides in the LH2 complex from *Mch. purpuratum*. The His and Asp residues that coordinate B828 and B800 respectively are highlighted in red. Residues with no sequence conservation are highlighted in green. The amino acids that were fitted into the final model are underlined.

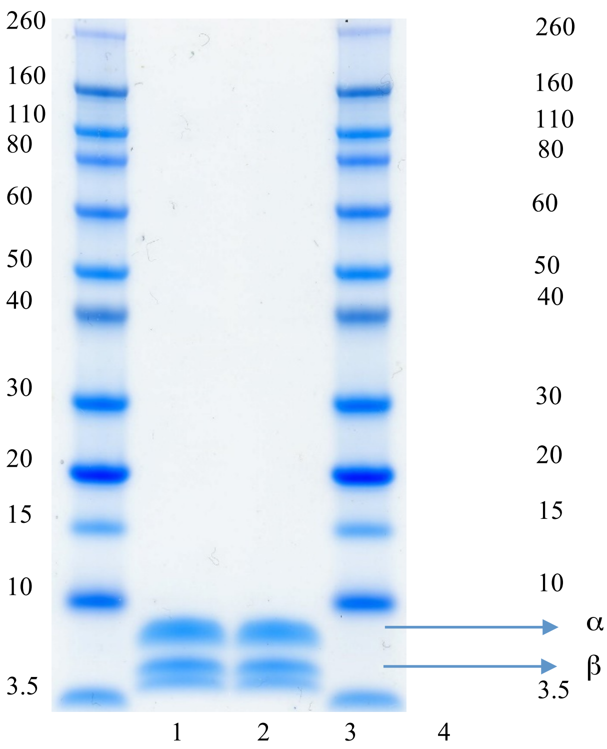


Fig. S2 **SDS-PAGE of purified LH2 from *Mch. Purpuratum***. Lanes 1 and 4: Protein ladder (in kDa). Lanes 2 and 3: Purified *Mch. pupuratum* LH2. A single α band corresponds to a mixture of 3 different α-polypeptides, and the 3 different β–polypeptides are separated into two bands. The pre-cast NuPAGE 4-12% Bis-Tris gel was run with NuPAGE MES SDS running buffer, then stained with Coomassie blue.

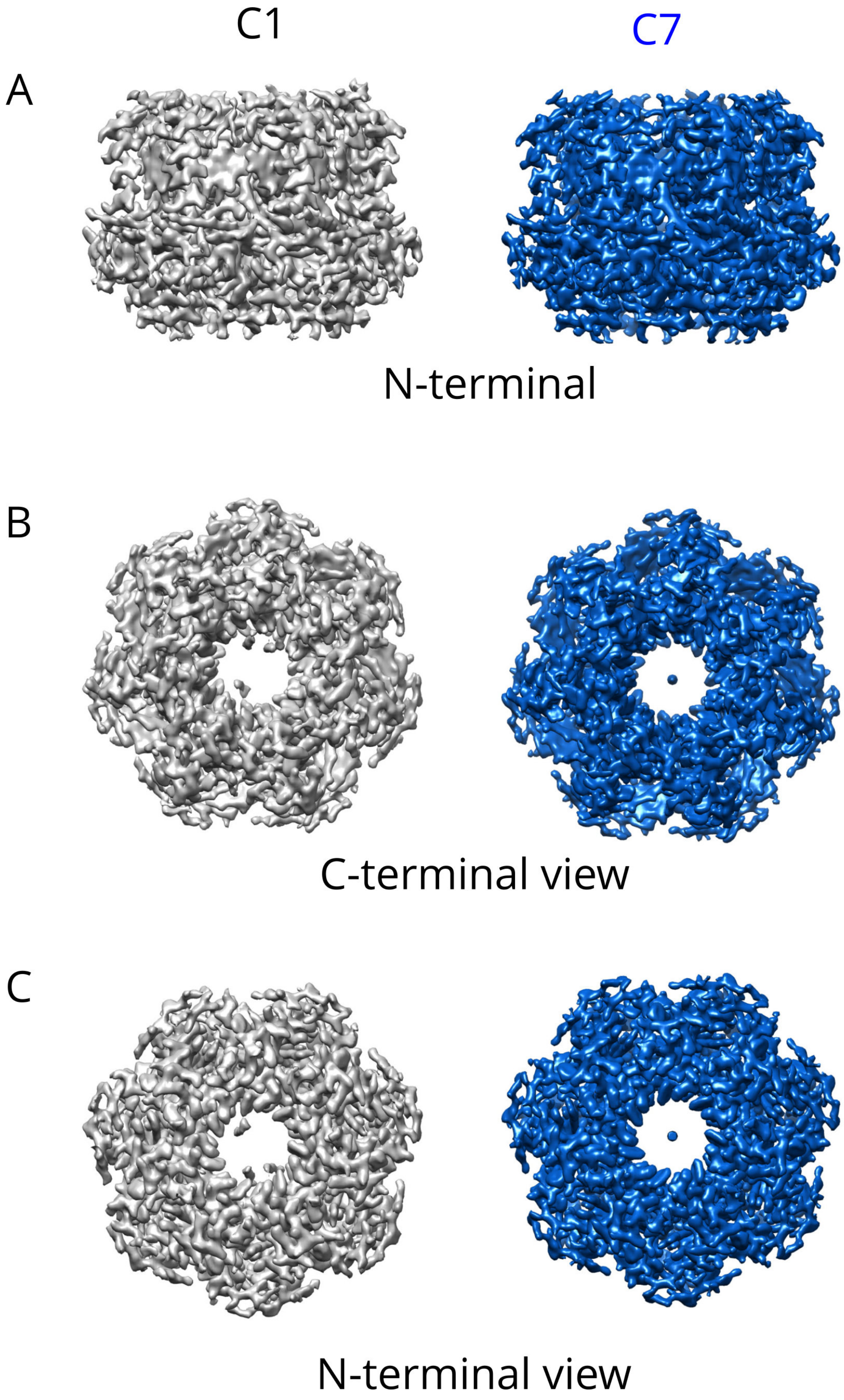


Fig. S3 **Comparison of C1 and C7 maps of the LH2 from *Mch. purpuratum*. (A)** Side view parallel to membrane level. The C1 map is shown in grey, and the C7 map in blue. (**B**) View from periplasmic side. (**C**) View from cytoplasmic side. For comparison, both C1 and C7 map volumes are adjusted to be equal.

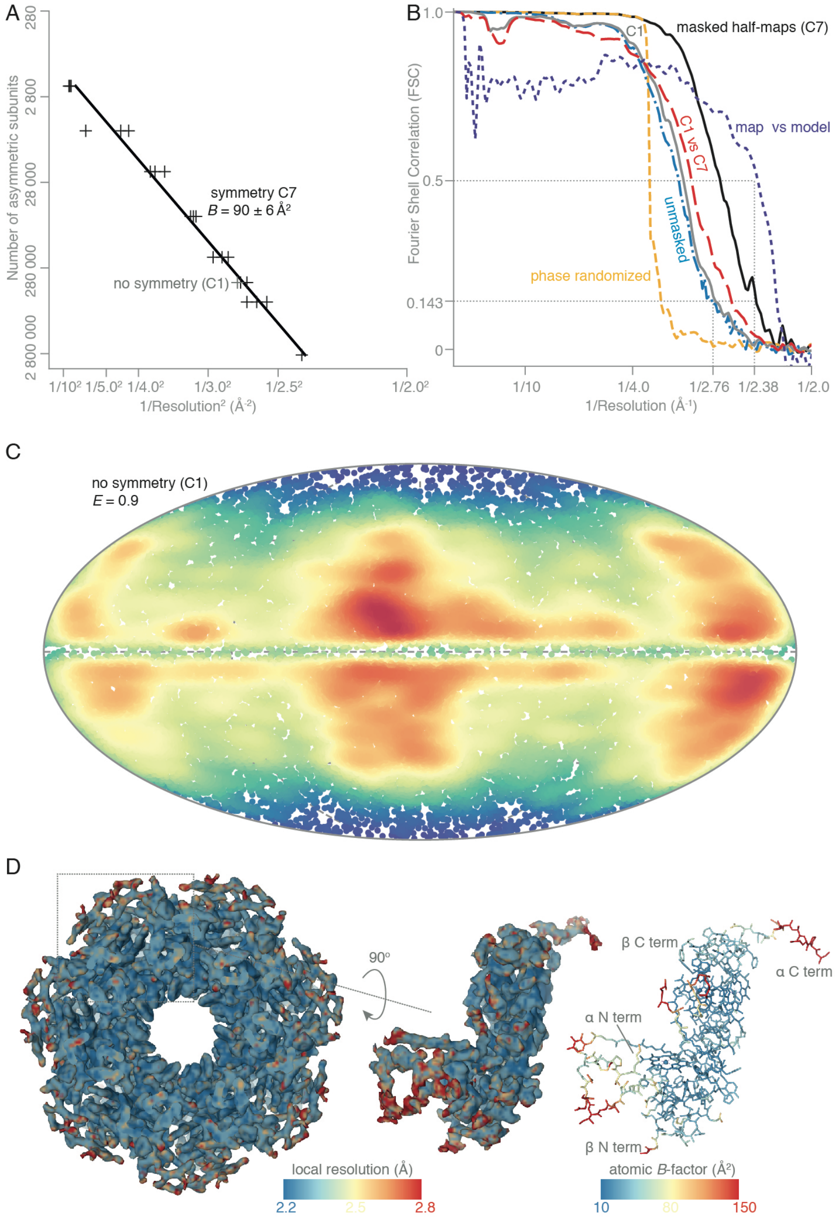


**Fig. S4 Characterization of the quality of the LH2 reconstruction**
(**A**) The number of asymmetric subunits required to reach a given resolution for the C7-symmetric LH2 is shown on a Rosenthal plot[36]. The *black line* is a linear fit to the resolutions from individual refinements on particle subsets (*black markers*), and corresponds to a B-factor of 90 ± 6 Å$^2$. The resolution of the refinement with all particles without symmetry is shown with the *grey marker*. It is the same, within error, as the resolution from one seventh of all particles with C7 symmetry imposed. (**B**) Fourier shell correlation (FSC) curves of the masked (*black solid line*), unmasked (*blue dash-dotted line*), and phase randomized (*yellow dashed line*) half-maps with

C7 symmetry, the masked half-maps without symmetry (C1, *grey solid line*), the C1 vs C7-symmetric masked half-maps (*red dashed line*), and the final zero dose extrapolated map, masked around the LH2 density only, versus the atomic model (*purple dashed line*) are shown, (**C**) The orientation distribution of a random subset of 10,000 LH2 particles, as determined from the refinement without symmetry, is shown on a Mollweide projection. Every point indicates a particle orientation, and the color scale corresponds to the probability distribution function of the orientations, ranging from 0 (*blue*) to $6 \times 10^{-5}$ (*red*). The efficiency[37] $E$ of the orientation distribution is 0.9, indicating nearly uniform Fourier space coverage. (**D**) The final, C7-symmetrised map is colored by local resolution, as determined in ResMap[38]. The local resolution ranges from 2.2 Å to 2.8 Å. An orthogonal view of one subunit of the heptamer, colored in the same way, is shown. The atomic model of the subunit is shown is the same orientation, colored by atomic $B$-factor, ranging from 10 Å$^2$ to 150 Å$^2$.

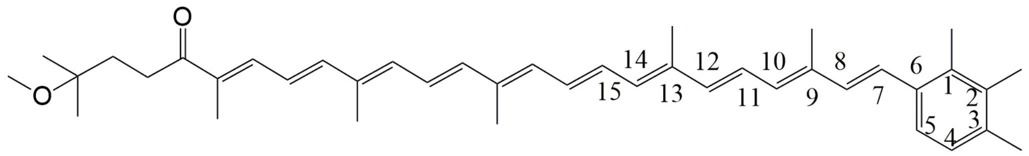


All-trans Okenone

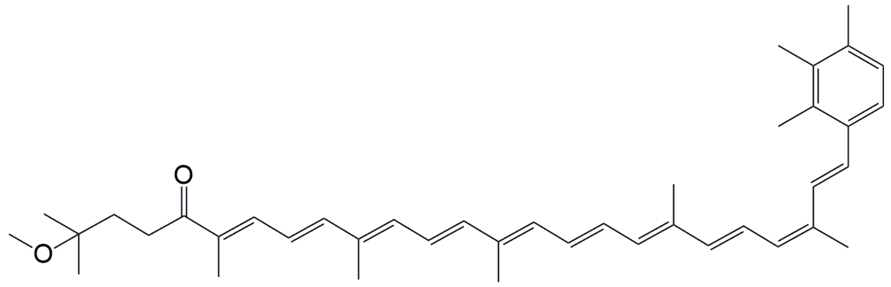


9-cis Okenone

**Fig. S5 Chemical structure of all-*trans* and 9-*cis* okenone**.

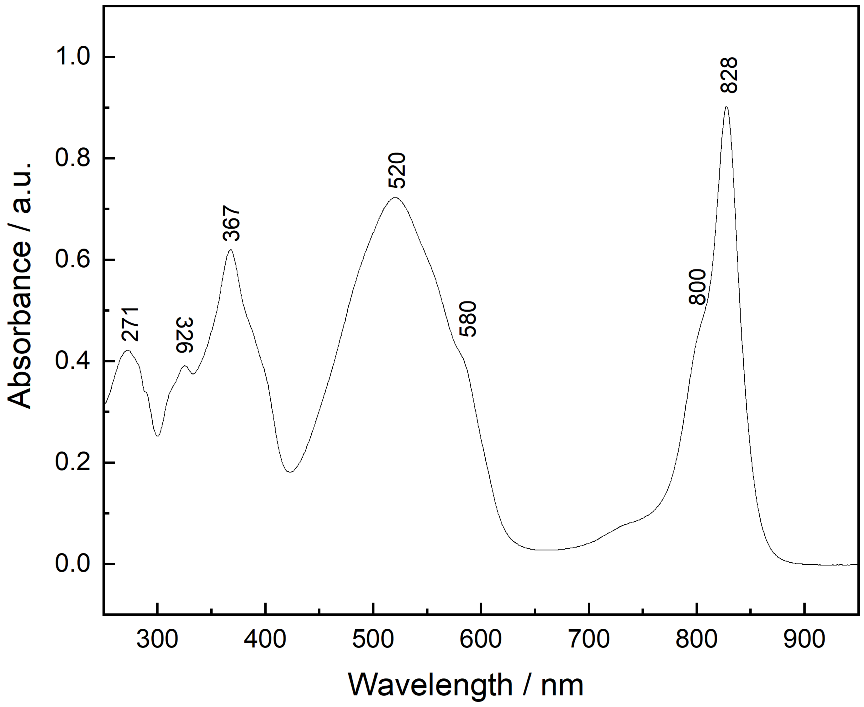


**Fig. S6 Absorbance spectrum of the purified LH2 complex from *Mch. Purpuratum***

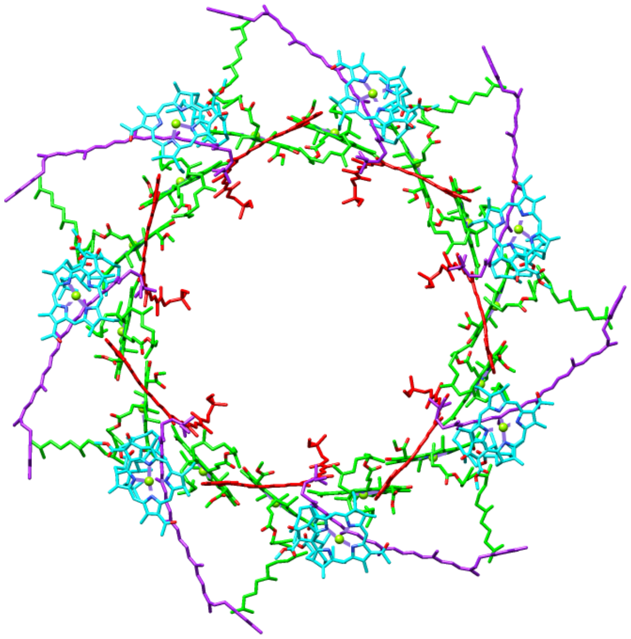

**Fig. S7 Pigment molecular organization in the LH2 complex from *Mch. purpuratum.*** The LH2 molecule is viewed from them cytoplasmic side. Color coding is the same as in Fig. 1

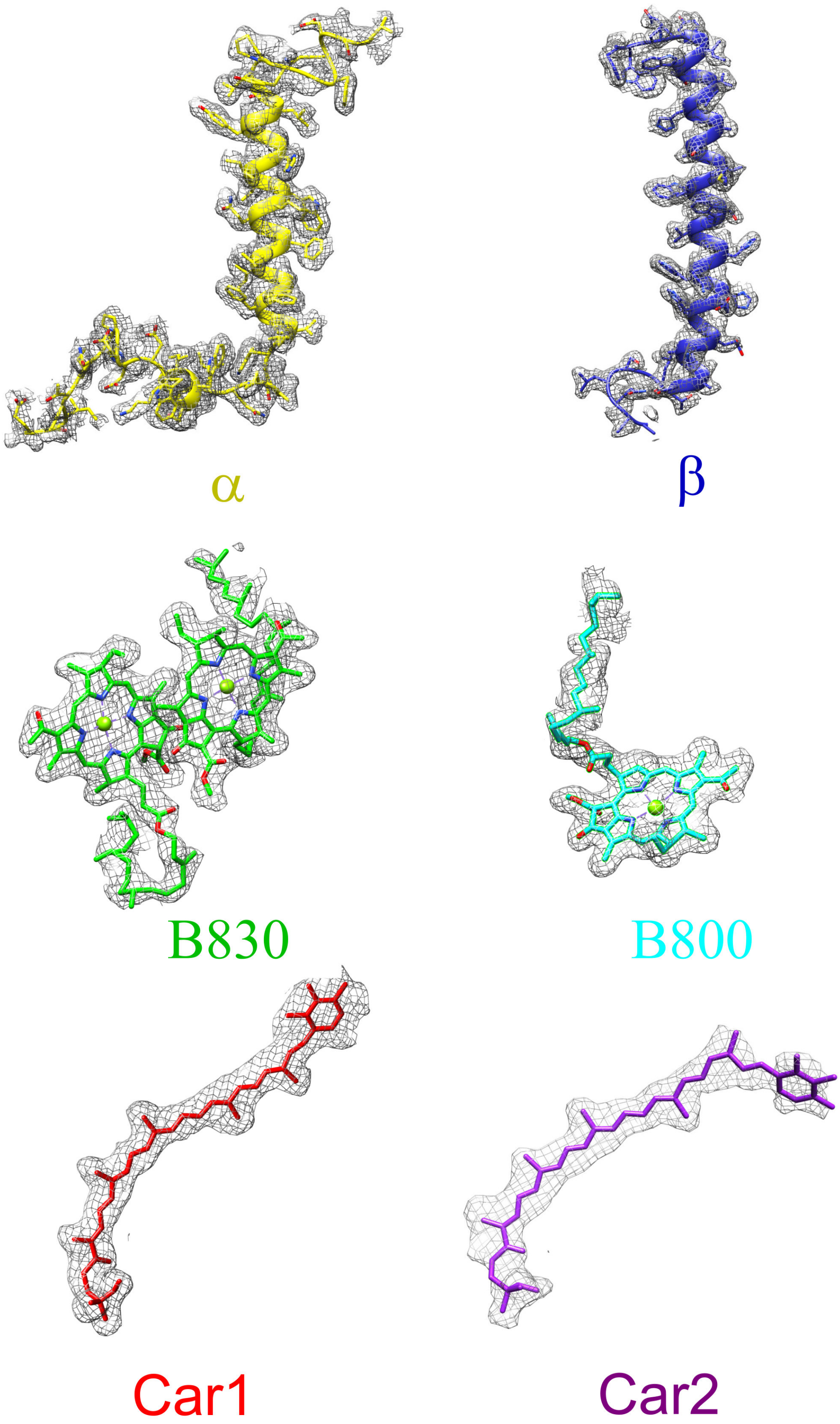


**Fig. S8 Cryo-EM density maps with models of proteins and pigment molecules in the LH2 complex of *Mch. purpuratum*.** The colour code is the same as in Fig. 1. The contour levels of the density maps were adjusted to just cover proteins and pigment molecules for clarity.

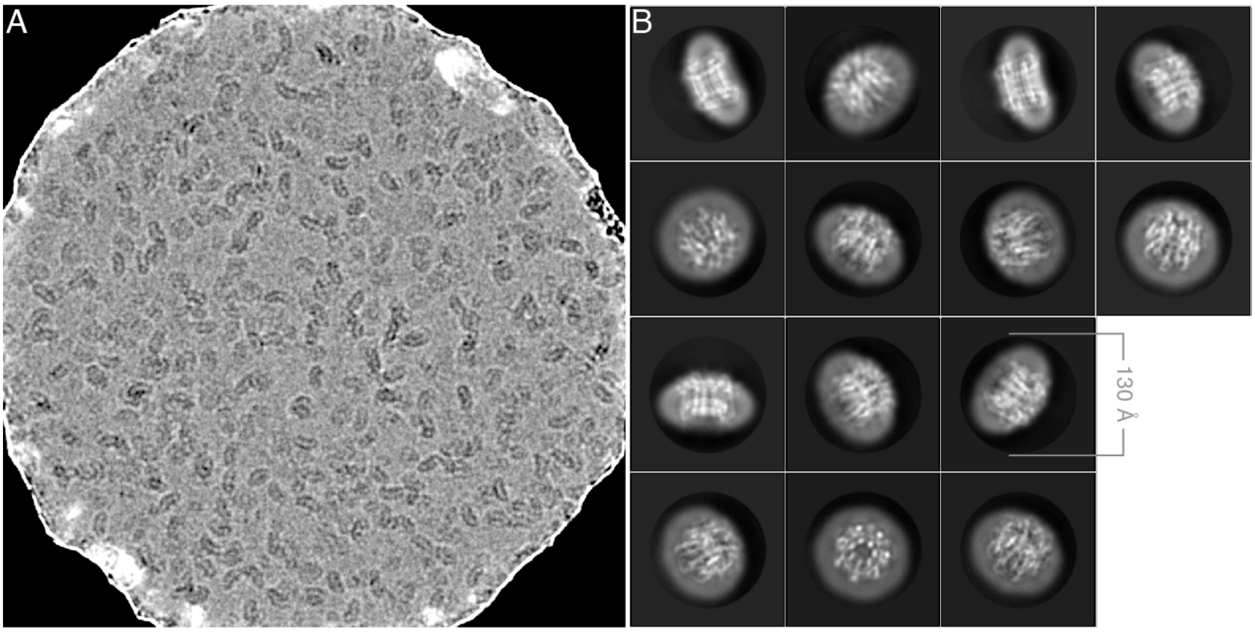


**Fig. S9 Electron cryomicroscopy of the LH2 complex from *Mch. purpuratum* on HexAuFoil specimen supports**. (**A**) A representative micrograph of LHC2 in vitreous ice in the 270 nm hole of a HexAuFoil grid at 2 μm defocus. The micrograph is low-pass filtered to 9 Å and the contrast is adjusted to improve the visibility of the particles. (**B**) These selected masked 2D class averages were used for the final reconstruction.

**Table S1: Cryo-EM data acquisition, model refinement and validation statistics.**

| **Protein source** | Photosynthetic bacterium |
|---|---|
| **Data collection and processing** | |
| Microscope | ThermoFisher Titan Krios G3i |
| Voltage (kV) | 300 |
| Camera | Falcon 4 |
| Energy filter | No |
| Energy filter slit width | N/A |
| Magnification | 120,000 |
| Defocus range (μm) | -0.4 to -3.0 |
| Mean defoucs (μm) | -1.8 |
| Pixel size (Å) | 0.646 |
| Electron flux ($e^-$/ $Å^2$/s) | 3.58 |
| Electron fluence ($e^-$/ $Å^2$) | 43.7 |
| Exposure time (sec/frame) | 0.29 |
| Electron fluence per frame (e/ $Å^2$/frame) | 1.04 |
| Number of frames per movie | 42 |

| | |
|---|---|
| Number of movies used | 9543 |
| Initial no. particle images | 1,330,154 |
| Final no. particle images | 414,511 |
| Estimated accuracy of translations (Å) (RELION) | 0.32 |
| Estimated accuracy of rotations($^{o}$) (RELION) | 0.98 |
| Symmetry imposed | C7 |
| Local resolution range | 2.2 to 2.8 |
| Map resolution (Å, FSC=0.143) | 2.38 |
| Resolution of unmasked reconstruction (Å, FSC=0.143) | 2.76 |
| Resolution of masked reconstruction (Å, FSC=0.143) | 2.38 |
| Specimen temperature | ~80K |
| Particle box size | $(512\ px)^2$ |
| **Refinement and validation** | |
| Refinement package | COOT, Refmac 5, PHENIX |
| Initial model | PDB 1LGH |
| Model resolution (Å, FSC=0.5) | 2.38 |
| Map sharpening B factor ($Å^2$) | -88 |
| **Model composition** | |
| Non-hydrogen atoms | 8442 |
| Protein residues | 791 |
| Molecular weight (kD) | 110.77 |
| Protein B factor ($Å^2$) | 20.55 |
| **RMS deviations** | |
| Bond length (Å) | 0.008 |
| Bond angle ($^{0}$) | 1.309 |
| **Validation** | |
| MolProbity | 2.66 |
| Clashscore | 9.45 |
| Poor rotamers (%) | 5.21 |
| EMRinger score | 5.13 |
| Cb deviations (%) | 0.00 |
| CaBLAM outliers (%) | 9.52 |
| **Ramachandran plot** | |
| Favoured (%) | 87.16 |
| Allowed (%) | 12.84 |
| Disallowed (%) | 0 |
| **PDB ID** | 6ZXA |
| **EMDB ID** | EMD-11516 |
